# Single-molecule visualization of twin-supercoiled domains generated during transcription

**DOI:** 10.1101/2023.08.25.554779

**Authors:** Richard Janissen, Roman Barth, Minco Polinder, Jaco van der Torre, Cees Dekker

**Affiliations:** Department of Bionanoscience, Kavli Institute of Nanoscience Delft, Delft University of Technology, Delft, South-Holland, 2629HZ, The Netherlands

## Abstract

Transcription-coupled supercoiling of DNA is a key factor in chromosome compaction and the regulation of genetic processes in all domains of life. It has become common knowledge that, during transcription, the DNA-dependent RNA polymerase (RNAP) induces positive supercoiling ahead of it (downstream) and negative supercoils in its wake (upstream), as rotation of RNAP around the DNA axis upon tracking its helical groove gets constrained due to drag on its RNA transcript. Here, we experimentally validate this so-called twin-supercoiled-domain model with *in vitro* real-time visualization at the single-molecule scale. Upon binding to the promoter site on a supercoiled DNA molecule, RNAP merges all DNA supercoils into one large pinned plectoneme with RNAP residing at its apex. Transcription by RNAP in real time demonstrates that up- and downstream supercoils are generated simultaneously and in equal portions, in agreement with the twin-supercoiled-domain model. Experiments carried out in the presence of RNases A and H, revealed that an additional viscous drag of the RNA transcript is not necessary for the RNAP to induce supercoils. The latter results contrast the current consensus and simulations on the origin of the twin-supercoiled domains, pointing at an additional mechanistic cause underlying supercoil generation by RNAP in transcription.

## INTRODUCTION

DNA supercoiling, the additional twisting of the double helix that leads to extended intertwined DNA structures, occurs in chromosomes in all domains of life. DNA supercoils contribute to the tight packaging of the genome and the control of essential genetic processes, including DNA replication, recombination, chromosome segregation, as well as regulation of the expression of genes [1]–[9]. Genomic DNA is subjected to severe torsional stress throughout the cellular life cycle due to transcription and replication [4], [10], [11] as well as due to the wrapping around proteins such as eukaryotic nucleosomes [12], [13] and nucleoid-associated proteins (NAPs) in prokaryotes [14], [15]. The built-up torsional stress in the DNA is relieved by the formation of supercoils that generate local intertwined loops (plectonemes) and by changes in DNA twist generated by ATP-dependent topoisomerases [1], [16], [17].

The fundamental cellular process of transcription, carried out by DNA-dependent RNA polymerases (RNAP) [10], [18] is a major source of supercoiling. It has become universally accepted that, during transcription, RNAP generates positive, right-handed supercoiling ahead (downstream) and compensatory negative, left-handed supercoiling in its wake (upstream), a phenomenon termed the *twin-supercoiled-domain model* [18]–[20]. Various experimental studies have provided strong evidence for this model. Bulk plasmid studies verified the induction of supercoils indirectly *via* the use of topoisomerases [19], [21], [22], while single-molecule studies showed the induction of supercoils of one handedness only individually [10], [23]. A direct experimental validation of twin-supercoiled-domain model with the implied simultaneous generation of up- and downstream supercoils has however still been lacking. Such a study would verify the model and address open questions such as the symmetry in the plectonemes sizes that are simultaneously generated up- and downstream or the necessity of an RNA transcript to generate enough rotational drag for supercoil induction.

Here, we address these questions at the single-molecule level using an established fluorescence-based technique with an array of stretched DNA molecules that allows the direct visualization and quantification of plectoneme diffusion and size, as well as of RNAP proteins over time [24], [25]. Our results demonstrate that promoter-bound RNAP first consolidates diffusive DNA supercoils into a single pinned plectoneme where it resides at the apical loop. Upon transcription along a torsionally constrained DNA template, we observed supercoils up- and downstream of the RNAP that were equally partitioned with respect to plectoneme size, an observation that confirms the twin-supercoiled-domain model. Intriguingly, experiments in the presence RNase A and H – which digest the RNA transcript and potential R-loops, respectively – showed that the viscous drag of the RNA transcript is not necessary for the RNAP to induce supercoils, and accordingly the bare RNAP can induce twist on its own, even in the low-viscosity aqueous environment of our in vitro experiments. This result contrasts the common explanation for the generation of the twin-supercoiled domains as well as data from simulations of the mechanics of transcription underlying the twin-supercoiled-domain model.

## MATERIAL AND METHODS

### Purification and labeling of *E. coli* RNAP and σ^70^

Wild-type *E. coli* RNA polymerase (RNAP) core-enzyme (α2ββ′ω) with a SNAP-tag at ß’ and the transcription initiation factor σ^70^ were expressed and purified as described previously [26]. Alexa647-O^6^-benzylguanine was attached to purified RNAP-SNAP according to the protocol provided by the supplier (New England Biolabs) and filtered using a Sepharose 6 gel-filtration column (Cytiva). The labeling efficiency of the RNAP core enzyme was measured to be 89% (Nanodrop; Thermo Fischer). RNAP-Alexa647 holoenzyme formation with σ^70^ was performed by incubating RNAP-Alexa647 with σ^70^ in a 1:5 ratio for 20 min at 30°C in a buffer containing 10 mM TRIS, 100 mM KCl, pH 7.9, and stored in 10 mM Tris-HCl, pH 7.9, 100 mM KCl, 0.1 mM EDTA, 0,1 mM DTT, and 50 % glycerol at −80°C.

### Synthesis of DNA constructs

Plasmids used for the construction of torsionally constrained 21kb-T7A1 and 38kb-T7A1 DNA constructs were previously described in detail [27]. The torsionally constrained 31kb-T7A1 DNA construct was made from plasmid #140-pBS-T7A1. This plasmid was made using a BsaI-HFv2 golden gate cloning reaction with destination plasmid #64, and donor plasmids #66, #133, #69, #65 and #71 (NEB E1601), as previously described [27], except plasmid #133-pGGA-T7A1. Plasmid #133-pGGA-T7A1 was made in several cloning steps: first plasmid #125-pGGA-T7A1rev was constructed using blunt ligation cloning. The backbone and insert fragment were made with Q5 DNA polymerase PCR (M0491, NEB), where the insert PCR fragments were made using primers JT418 (TTAAAATTTATCAAAAAGAGTATTGACTTAAAGTCTAACCTATAGGATACTTACAG) and JT406 (TAAATCTAACAAAATTCTATCCTGGGACATGCACTCTAGTCAGGATGATGGTGATG) on plasmid #92-pSC-T7A1reverse [27]. The backbone fragment was a PCR fragment using primers NH1 (AAAGAAATCAAAGGCGCGGAC) and NH2 (CACGATCCCGTTTTGTGAGTTG) on #67-pGGA-JT294JT295 [27]. Before the ligation reaction, the insert PCR fragment was phosphorylated with P4 polynucleotide Kinase (M0201, New England Biolabs, UK) and the destination PCR-fragment was DPN1 treated (R0176, New England Biolabs, UK). These two fragments where ligated together using T4 DNA ligase (M0202, NEB). The resulting plasmid can have the T7A1 promoter in two different orientations, upon analysis we only got plasmids in which the T7A1 was orientated towards the NH2-primer, creating plasmid #125-pGGA-T7A1rev. To have the T7A1 orientated towards the other side in the end plasmid, we swapped the BsaI overhangs in this plasmid. For this we performed two subsequent KLD cloning reactions (M0554, New England Biolabs, UK). In the first reaction we made a PCR-fragment with Q5 DNA polymerase (M0491, New England Biolabs, UK) using primers JT403 (CCTGTAGTCTTCTTAATTAAGACGTCAG) and JT447(CGACAAGGTCTCCAGCCCGCGATGGTTGGAGTTCCAG) on plasmid #125, creating plasmid #131. Subsequently, plasmid #133-pGGA-T7A1 was made in a second reaction conducting a KLD reaction using PCR with primers JT401 (GTACCAAGTCTTCGAATTCGGATC) and JT446(CGACAAGGTCTCCATACGCGACTTACCATGTAT) on plasmid #131.

Linear DNA constructs were prepared using freshly isolated #92-pSC-T7A1reverse (21kb-T7A1), #140-pBS-T7A1(31kb-T7A1), and #189-pBS-T7A1-ParS(38kb-T7A1) midiprep DNA, using Qiafilter plasmid midi kit (Qiagen, Germany). These plasmids were digested with NotI-HF and XhoI for 2 h at 37°C (R3189, R0146, New England Biolabs, UK) and heat-inactivated for 20 min at 80°C.

For all the constrained fragments we created handles – additional pieces of DNA that were attached to the linear DNA constructs – containing multiple biotin by a PCR of pBluescript SK+ (for 21kb-T7A1 and 31kb-T7A1 constructs) or of #186-pBS-250handle (for 38kb-T7A1 construct) using primers CD21(GACCGAGATAGGGTTGAGTG) and CD22(CAGGGTCGGAACAGGAGAGC). We used a ratio of 1:5 of Biotin-16-dUTP (NU-803-BIO16, JenaBioscience, Germany):UTP in the PCR reaction mix during synthesis of a 1,246 bp, multi-biotin containing PCR-fragment. The handles where split in two fragments by digesting with XhoI or NotI-HF; after PCR and digestion reactions, the handles were purified using a PCR clean up kit (A9282, Promega).

Finally, we mixed the digested DNA constructs and handles in a 1:10 molar ratio and ligated them together using T4 DNA ligase (M0202, New England Biolabs, UK) at 16°C overnight, which was subsequently heat-inactivated for 10 min at 65°C. The 38kb-T7A1 constructs were, before cleanup, digested for 1h at 37°C with SrfI (R0629) and then heat-inactivated for 20 min at 65°C. We subsequently purified all resulting DNA constructs from the access of handles and other DNA fragments using an ÄKTA Start (Cytiva), with a homemade gel filtration column containing 46 ml of Sephacryl S-1000 SF gel filtration media, run with TE + 150 mM NaCl buffer at 0.5 ml/min.

### Dual-color HiLo fluorescence microscopy

Details of the experimental setup were described previously [25], [28]. Briefly, an Olympus IX81 TIR microscope, equipped with a 60x oil-immersion objective (CFI APO TIRF, NA 1.49, Nikon), two lasers (532 nm and 640 nm; Cobolt), and a sCMOS camera (PrimeBSI; Teledyne Photometrics), was used to image RNAP-Alexa647, and DNA molecules *via* Sytox Orange (Thermo Fischer) intercalating dyes. Fluorescent emission was first filtered by a dichroic mirror (FF635-Di02; Semrock) and for the Sytox Orange and Alexa647 channels the band-pass filters FF01-731/137 (Semrock) and FF01-571/72 (Semrock) were used, respectively.

### Single-molecule flow cell preparation

The flow cells used in this study have been described in detail previously [25], [28], [29], with the addition that the surface were PEGylated four times. In short, a 24×60 mm quartz glass slide and a coverslip (#1.5, Menzel GmbH, Germany) of equal size were coated with polyethlylene glycol (PEG) to suppress nonspecific binding of biological material and Sytox Orange. 2% of the PEG molecules were biotinylated for the DNA immobilization via biotin−streptavidin linkage. The quartz slide and coverslip were sandwiched with strips of double-sided tape at a distance of 5 mm between them, forming shallow sample channels. Two holes in the quartz glass slide serve as the inlet and outlet of the flow. Pipet tips serve as reservoir at the inlet and tubing was connected at the outlet and at a syringe to induce flow using a syringe pump. Typically, a flow channel holds 10 μl of solution.

### Single-molecule supercoiling assay

DNA supercoiling was induced by the addition of the DNA intercalator Sytox Orange to a 31 kpb DNA construct, as described previously [24]. To induce negative supercoiling, 250 nM Sytox Orange was used during flushing the DNA constructs into the flow cells until they are immobilized with both ends on the surface. Subsequently, the Sytox Orange concentration was reduced to 50 nM which resulted in negative supercoiling. To induce positive supercoiling, 30 nM Sytox Orange was used during flushing the DNA constructs into the flow cells. An increase of Sytox Orange afterwards to 200 nM resulted in positive supercoiling. The experiments probing the ability of bound RNAP (in open complex conformation) to pin plectonemes were conducted in the same manner, following prior RNAP binding and stalling to the torsionally constrained DNA construct. All measurements were performed in an imaging buffer (40 mM Tris HCl, 50 mM KCl, 5 mM MgCl_2_, 2 mM Trolox, 30 mM Glucose, 0.1 mM DTT, 10 nM Catalase, 18.75 nM Glucose Oxidase, 0.25 µg/ml BSA) at room temperature with an acquisition frequency of 0.2 or 0.5 Hz. The two-color fluorescence emission was imaged in series with a 100 ms exposure time for each wavelength.

### Formation of stalled transcription elongation complexes

Prior to anchoring the DNA construct onto the streptavidin-functionalized flow cell surface, RNAP-Alexa647 holoenzyme was stalled at the position A29 nt after the T7A1 promotor as previously described [30], [31]. To do so, 6 nM of RNAP holoenzyme was added to 3 nM DNA construct in stalling buffer (20 mM Tris, 150 mM potassium glutamate, 10 mM MgCl_2_, 1 mM DTT, 40 µg/ml BSA, pH 7.9) and incubated for 10 min at 30°C. Afterwards, 2.5 µM ATP, CTP, GTP (GE Healthcare Europe), and 100 µM ApU (IBA Lifesciences GmbH) were added to the solution and incubated for additional 10 min at 37°C. To ensure measuring the transcription dynamics of single RNAP ternary complexes, a final concentration of 100 µg/ml heparin (Sigma) was added and incubated for 10 min at 30°C to sequester free RNAP and those that were weakly associated with the DNA.

### Single-molecule transcription assay

The RNAP-Alexa647:DNA ternary complex was diluted to a final concentration of 100 pM in imaging buffer (40 mM Tris HCl, 50 mM KCl, 7 mM MgCl_2_, 2 mM Trolox, 30 mM Glucose, 0.1 mM DTT, 10 nM Catalase, 18.75 nM Glucose Oxidase, 0.25 µg/ml BSA, 100 nM Sytox Orange) before flushing into the flow cell at the speed of 6 µl/min; unbound molecules were washed out with imaging buffer. One minute after starting the imaging by alternate excitation with 100 ms exposure times for DNA-Sytox Orange (561 nm laser), and RNAP-Alexa647 (647 nm laser) followed by an 800-1,800 ms pause before the next frame, a final concentration of 1 mM of all four rNTPs (GE Healthcare Europe) was added to re-initiate transcription and being recorded for 5 min.

RNase A (100 nM; Quiagen) or RNase H (0.5 U/µl; Thermo Fischer) were afterwards added in imaging buffer complemented with 1 mM rNTPs, and recorded for another 5 min. Afterwards, to verify if the imaged DNA tethers remained torsionally constrained, 1 µM Sytox Orange in imaging buffer was added and recorded for another 3 min. All measurements were performed at room temperature with an acquisition frequency of 1 or 2 Hz.

### Data analysis

From the recorded fluorescence image stacks, areas of single DNA molecules were cropped from the field of view and analyzed separately with custom-written scripts [32] in python 3.8 and Igor Pro V6.37. To reduce image noise, the cropped DNA images were smoothed with a median filter with a window size of 3 pixels and background subtraction was performed using the “white_tophat” operation provided in the *scipy* python module. Kymographs were then constructed along the long axis of the stretched DNA construct.

The DNA ends were determined by a *peak peeling* algorithm [33], where several frames prior the induction of supercoils or addition of rNTPs were temporally averaged to gain a fluorescence intensity profile of the DNA molecule. A series of Gaussian peaks with a full-width-half-maximum of the microscope’s point spread function were placed at – and subtracted from - the maximum DNA profile intensities. This procedure was repeated until <10% of original integrated area of the DNA intensity profile remained. The location of the most outer Gaussian peaks then corresponded to the DNA ends.

For each image frame, high intensity foci corresponding to DNA plectonemes or RNAP were identified and tracked using the *scipy.find_peaks* and *trackpy* algorithms as previously described [33], as well as the calculation of the base pair size of each detected plectoneme that existed for a minimum of three consecutive frames. Pixel positions of the detected plectonemes and RNAP were converted to micrometer and genomic positions considering a measured pixel size of 109 nm/px and the DNA end-to-end length of known base pair length of the used DNA constructs.

All statistical analyses in this study consisted of unpaired, two-tailed t-tests.

## RESULTS AND DISCUSSION

### RNAP promoter binding pins supercoils into a single plectoneme

First, we examined the DNA topology that resulted upon binding of RNA polymerase (RNAP) onto a supercoiled DNA molecule. Recent single-molecule and molecular-dynamics-simulation studies of sequence-dependent bending of linear DNA demonstrated that a high local DNA curvature poses a key factor that facilitates the pinning of a plectonemic supercoil [25], [34]. These results suggest that DNA-binding proteins that induce a strong DNA bending may also pin genomic supercoils, independent from the intrinsic curvature of the DNA sequence. Supporting this notion, various nucleoproteins from all three domains of life, that bridge, bend, or wrap DNA around them, have shown the ability to stabilize the plectoneme topology [1], [12], [35]–[37]. While many nucleoproteins mostly serve an architectural role, the abundant DNA-dependent RNAPs bind, transcribe, and dissociate from the genomic DNA with high rates during the entire cellular life cycle [38]–[40]. As the DNA is both locally melt and strongly bent by ~90-100° by RNAP (RNAP_OC_) upon open complex formation [41]–[44], we reasoned that bound RNAP_OC_ might strongly pin plectonemes. Consequently, we expected that binding of RNAP onto a supercoiled DNA molecule would yield a pinned plectonemic supercoil at that position, with abound RNAP_OC_ residing at the apex of the plectoneme.

We sought out to verify this hypothesis at the single-molecule level using a multiplexed fluorescence method involving intercalation-induced supercoiling, which tracks the formation, size, and diffusion behavior of single plectonemes over time [24], [25]. The assay consisted of a torsionally constrained 21 kbp DNA molecule with one T7A1 promoter that is localized off center (~8 kbp from one DNA end) and multiple biotins at both DNA end regions to constrain its twist (Figure 1A). The DNA molecules were exposed to 250 nM Sytox Orange (SxO) intercalating dye and introduced into a sample chamber coated with streptavidin at constant flow, yielding DNA molecules that were tethered with both ends to the surface. Negative supercoiling was subsequently achieved by reducing the SxO concentration to 50 nM (see Materials and Methods), as a reduction of the amount of intercalators reduces the positive twist in the DNA, leading to negative plectonemes. The latter were readily observed as dynamic puncta of high fluorescence intensity that were diffusing along the DNA construct (Figure 1B; Supplementary Video V1), which represent a known visual signature of plectonemes [24], [25], [45]. Analogously, positive supercoils were induced by increasing the SxO concentration after tethering of the DNA molecules to the surface in a buffer with low SxO concentration (see Materials and Methods). In our assay, supercoils were thus induced by intercalating dyes without the need for direct mechanical manipulation of the DNA twist. Kymographs showed diffusing plectonemes as bright traces which, in agreement with previous observations [24], [25], [45], were highly dynamic in their nucleation, diffusion, and termination behavior (Figure 1B), as determined by tracking the position and quantifying the relative intensity of the plectoneme puncta in each frame (Figure 1C; Supplementary Figure 1).

**Figure 1.**
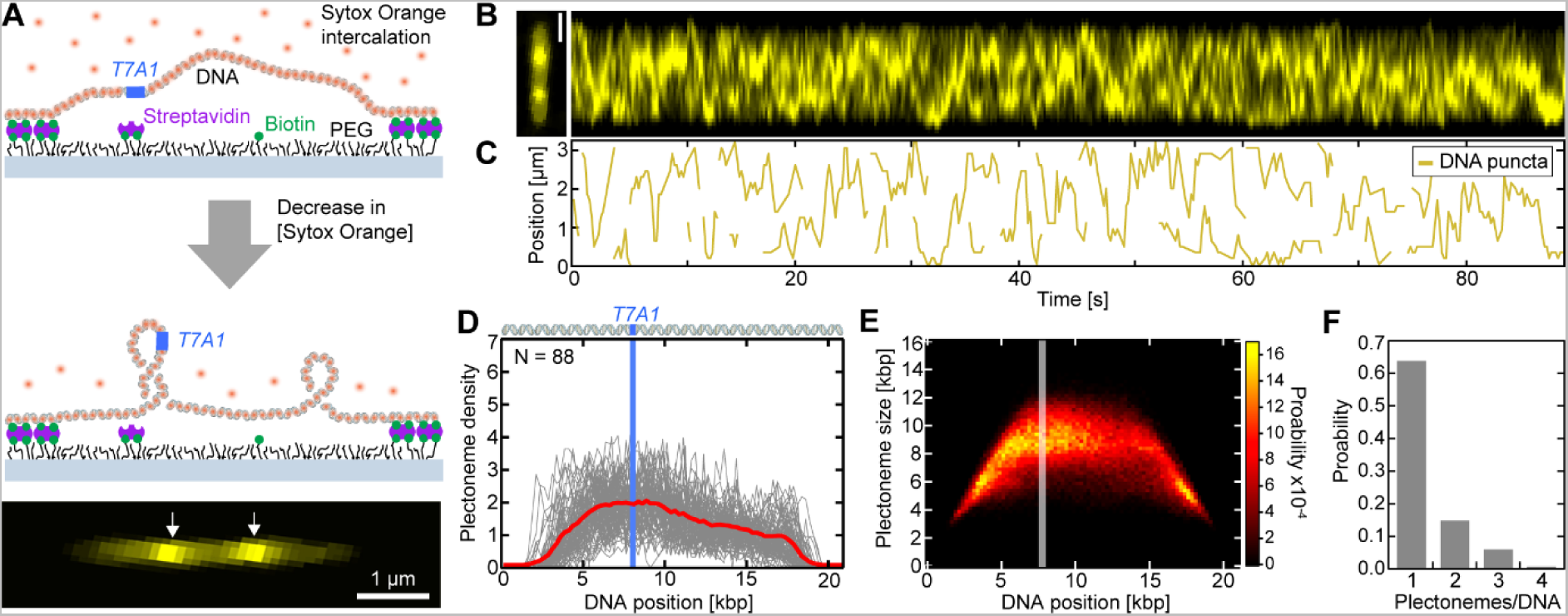
Visualization of plectonemes diffusing along negatively supercoiled DNA. (**A**) Schematic of the single-molecule supercoiling assay. (Top) A 21-kbp-long torsionally constrained DNA molecule with a T7A1 promoter site (blue) is flow-stretched and doubly-tethered onto a PEGylated surface via multiple streptavidin-biotin linkages at its end. (Center) The partial removal of the intercalating dye Sytox Orange (orange) after tethering of the DNA induces negative twist causing negative plectonemic supercoils into the DNA. (Bottom) Representative fluorescence image of a negatively supercoiled DNA. White arrows depict areas of higher DNA intensities that are signatures of plectonemes. (**B**) Fluorescence image (left) and intensity kymograph (right) of negatively supercoiled DNA. (**C**) Tracked positions of plectoneme puncta from (B) versus time, depicting nucleation, diffusion, and termination of plectonemes. (**D**) Plectoneme density versus DNA position for N = 88 individual DNA molecules (grey lines) and their average (red line). Top shows a schematic of the DNA molecule with the blue line indicating the position of the T7A1 promoter site. (**E**) Probability distribution of plectoneme size versus DNA position. T7A1 promoter position is depicted by the grey line. (**F**) Probability distribution of the number of plectonemes observed simultaneously, showing that multiple plectonemes can co-exist. Scale bars depict 1 µm. See also Supplementary Figure S1, Supplementary Figure S2 for positively supercoiled DNA, and Supplementary Video V1.

To quantitatively evaluate the DNA position dependence of individual plectonemes occurrence, we constructed a plectoneme density distribution (Figure 1D), which is the distribution of the genomic positions of individual plectonemes detected from all time frames pooled over N = 88 measured DNA molecules. The observed plectoneme density showed a fairly homogenous distribution across the 21 kbp DNA, with a slight increase in occurrence near the ±4 kbp region around the T7A1 promoter site. Proximate to the DNA molecule ends (<1.2 kbp and >19.8 kbp), the plectoneme density strongly decreased, which we attribute to the surface-attached handles in our assay (~600 bp at each DNA end; see Material and Methods) that limit the occurrence of sizeable plectonemes close to the DNA attachment sites. Similar to the DNA position-dependent probability distribution, the plectoneme size distribution (Figure 1E) showed a rather homogeneous distribution with a slight increase near the promotor site; we observed an average plectoneme size of ~8 kbp. An increase of the plectoneme localization probability at and around transcription start sites (TSSs) was proposed earlier from observations that the DNA curvature near bacterial TSS are relatively high [46]– [48], facilitating plectoneme nucleation, which was verified *in vitro* and *in silico* [25]. Figure 1F presents the probability distribution of the number of plectonemes that co-existed at each frame (exposure time 100 ms) for all DNA molecules measured. Most of the time (64% probability), a single plectoneme was present, while we observed multiple co-existing plectonemes, up to four plectonemes – in good agreement with previous observations on a similar DNA construct [24]. Notably, all these observations were independent of the supercoil handedness, as similar results were obtained for positively supercoiled DNA (Supplementary Figure S2A-D).

Subsequently, we extended the assay by adding a Alexa647-labeled *E. coli* RNAP which was stalled 29 nt after the T7A1 TSS (see Material and Methods). Here, a pre-stalled RNAP ternary complex with the 21 kbp DNA construct was introduced to the sample chamber in the presence of 250 nM SxO (Figure 2A). After DNA end-tethering to the surface, negative supercoiling was induced by the reduction of the SxO concentration to 50 nM. In contrast to bare DNA (Figure 1B), the DNA kymograph (Figure 2B) in the presence of stalled RNAP_OC_ strictly showed only one non-diffusing punctum that aligned with the RNAP position over the entire experiment time (Figure 2BC; Supplementary Video V2). This result directly showed that the RNAP_OC_ was able to efficiently pin all supercoils into a single plectoneme at the RNAP_OC_ stalling position. Our quantitative analysis over N = 79 DNA molecules with promoter-bound RNAP ternary complexes provides further support to this notion: the genomic position-dependent plectoneme density distribution (Figure 2D), as well as the plectoneme size probability distribution (Figure 2E) confirmed that plectonemes existed only near the T7A1 promoter site where RNAP was stalled. The presence of only one plectoneme (99% probability) detected in all experiments (Figure 2F) further indicated that supercoil structures were merged into the one pinned plectoneme of ~9 kbp size. Substantiating the above observations, the genomic plectoneme and RNAP_OC_ positions correlated fully at each frame for both supercoiling directions (Figure 2G). The observed plectoneme pinning by RNAP_OC_ was independent from the supercoil handedness, as the experiments with positively supercoiled DNA (Supplementary Figure S2F-H) provided identical results.

**Figure 2.**
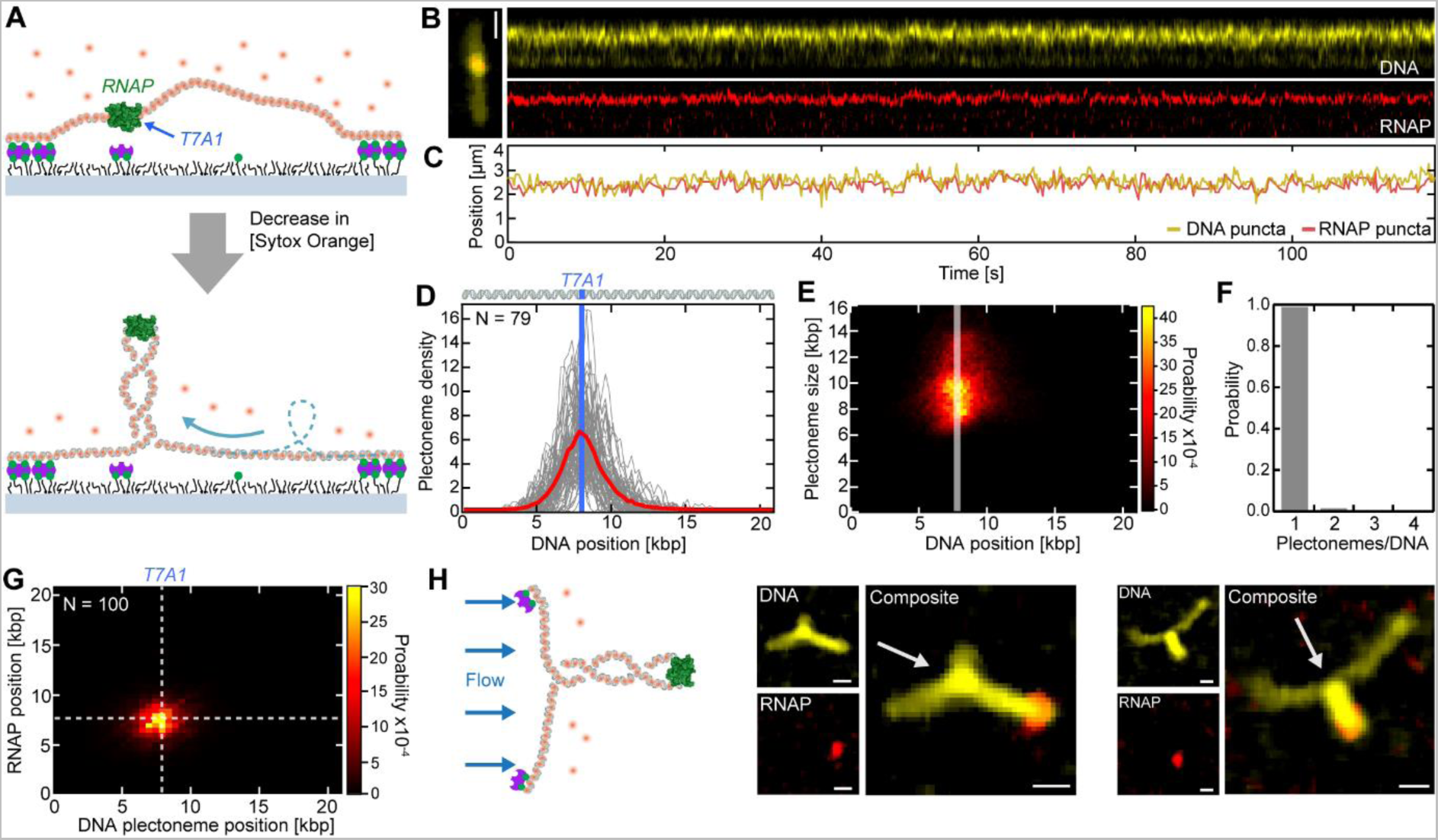
RNAP merges all DNA supercoils into one large pinned plectoneme. (**A**) Schematic of the assay with an RNAP (green) that was stalled at the T7A1 promoter site, yielding a merging of supercoils into one large plectoneme that is pinned in position by the RNAP residing at the plectoneme apex. (**B**) Representative fluorescence kymographs of DNA (top; yellow) and RNAP (bottom; red). (**C**) Tracked DNA (yellow) and RNAP (red) puncta positions on the DNA from (B), which clearly overlap during the entire experiment. (**D**) DNA position-dependent plectoneme densities in the presence of bound RNAP for N = 79 individual DNA molecules (grey lines) and their average (red line). T7A1 promoter position is depicted by the blue line. (**E**) DNA position-dependent plectoneme size probability distribution from (D). T7A1 promoter position is indicated by the grey line. (**F**) Probability distribution of the number of plectonemes co-existing simultaneously, showing only one plectoneme upon binding of RNAP to negatively supercoiled DNA. (**G**) Probability distribution of RNAP and DNA plectoneme positions for positively (N = 50) and negatively (N = 50) supercoiled DNA molecules. T7A1 position is displayed with white dashed lines. (**H**) Flow assay (left) and example dual-color fluorescence images (right) of 38-kbp-long torsionally constrained DNA construct under side-flow, demonstrating that RNAP was positioned at the apex of the plectonemes. White arrows depict flow direction. Scale bars depict 1 µm. See also Supplementary Figure S3, Supplementary Figure S2 for positively supercoiled DNA, and Supplementary Videos V2, V3.

We next probed the physical location of the stalled RNAP_OC_ along the plectoneme structure. To do so, we introduced a pre-stalled ternary RNAP complex using a torsionally constrained 38 kbp DNA construct with a T7A1 promoter site to the sample chamber, again in presence of 250 nM SxO, followed by a SxO concentration reduction to 50 nM after tethering to induce negative supercoiling. In contrast to the assays described above, we now applied a side flow perpendicular to the DNA orientation, to visualize the RNAP position relative to the plectoneme structure (see Figure 2H for two examples). For all measured DNA molecules (N = 10), we found that the stalled RNAP_OC_ was localized at the plectoneme apex (Supplementary Figure S3, Supplementary Video V3).

Our results that bound RNAP_OC_ is able to pin plectonemes and that RNAP resides at the plectoneme apical loop confirm early AFM and EM indications that were made in dry conditions for bacterial and eukaryotic RNA polymerases [49]–[51]. In the cellular environment, bound RNAP transcription-initiation complexes were also shown to preferentially localize at apical loops of DNA plectonemes [52], [53]. Our results are consistent with the notion that plectoneme pinning due to promoter-RNAP interactions represents a conserved architecture for transcription-initiation complexes in bacteria and eukaryotes [54].

### Transcribing RNAP generates supercoils with equal density up-and downstream

Transcription by RNAPs represents a major process in generating twist and supercoils in the cellular chromosome [4]. The currently accepted underlying mechanism was first postulated by Maaløe and Kjeldgaard in 1966 [55]. The authors suggested that while RNAP translocates along the helical groove of the duplex DNA, the viscous drag exerted by the RNAP and its nascent transcript and associated mRNA-processing factors hampers the RNAP from rotating around the DNA helix in the macromolecular crowded environment of the cell. Torsional anchoring of the transcriptome to other molecular structures in the cell or membrane, for example through membrane-bound ribosomes and transcription factors, can contribute as well to such a torsional constraining of the RNAP[56], [57]. Consequently, the twist generated by the processing transcriptome would not be accommodated by rotating the RNAP but instead be induced into the DNA [21], [58]–[60]. Since genomic DNA is torsionally constrained by cellular structures and architectural nucleoproteins (nucleosomes, NAPs, other DNA-binding proteins) [1], [4], [8], [13], [61], the translocation of RNAP during transcription causes overwinding of the DNA downstream from the RNAP, and compensatory underwinding upstream, resulting in, respectively, positive (right handed) and negative (left handed) supercoils. This so-called ‘twin-supercoiled-domain model’ of transcription has gained support from experimental *in vitro* and *in vivo* studies as well as from theoretical approaches over the past decades [10], [18]–[22], [58], [59], [62], [63]. Nonetheless, direct evidence of the simultaneous generation of up- and downstream supercoils as well as their symmetry in density remains to be provided experimentally.

We used our single-molecule visualization assay to measure this and validate the twin-supercoiled-domain model. We introduced the Alexa647-labeled RNAP again in the form of a stalled RNAP ternary complex using a torsionally constrained 31 kbp DNA construct with a T7A1 promoter site trailing a *rpoB* gene. To monitor the induction of supercoils during transcription, we kept the SxO concentration constant at 100 nM throughout the tethering of the DNA molecules to the surface and subsequent re-initiation of transcription by the addition of 1 mM of all four NTPs (Figure 3A). Any supercoils will thus not being induced by a change in SxO concentration but by the transcriptional activity of RNAP alone. By tracking the position and size of supercoils as well as the RNAP over time (Figure 3B), we observed that the transcribing RNAP generated supercoils located up- and downstream relative to the RNAP position after the addition of NTPs in our buffer (Figure 3C; Supplementary Video V4) – in direct support of the twin-supercoiled-domain model. We monitored notable fluctuations in the RNAP position along the DNA template, which we attribute to stochastic changes in plectoneme size and possibly nucleation/termination of plectonemes up- and downstream, as observed in any torsionally constrained DNA (cf. Figure 1; [24]). Supporting this notion was the fact that such position fluctuations were not detected during the transcription of a torsionally unconstrained, linear DNA template (Figure 1; Supplementary Figure S4; Supplementary Video V5).

**Figure 3.**
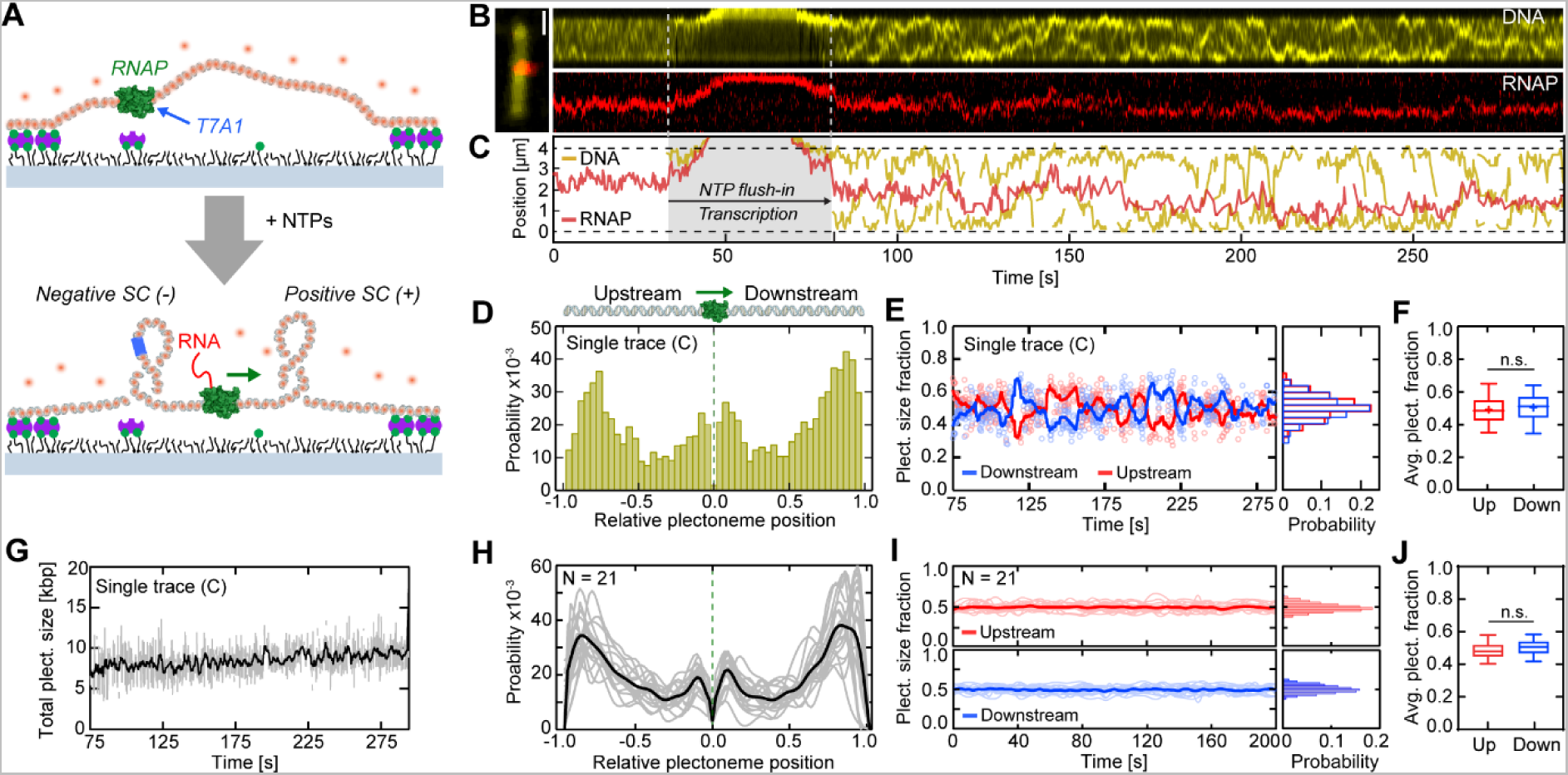
Transcription generates an equal portion of downstream and upstream supercoils. (**A**) Schematic of the transcription assay. Top: binding and stalling of an RNAP to a 31 kbp, torsionally constrained but not supercoiled DNA molecule at a T7A1 promoter site (blue). Bottom: restart of transcription by addition of NTPs generates downstream and upstream supercoils. (**B**) Example fluorescence kymographs of DNA (top, yellow) and RNAP (bottom, red). (**C**) Tracked DNA (yellow) and RNAP (red) puncta positions from (B) versus time, showing the absence of supercoils before NTP addition and supercoils up- and downstream of RNAP after the transcription restart. (**D**) Plectoneme probability distribution (from trace in (C)), relative to the RNAP position (position 0) and the DNA tether points (position ±1). (**E**) Fraction of up- and downstream plectoneme size (from trace in (C)) during transcription versus time. Right panel displays their probability distributions. Solid lines depict moving average-filtered data (1 Hz). (**F**) Average up- and downstream plectoneme size fractions from panel (E) showing no statistical difference. (**G**) The total size of supercoils (i.e., sum of upstream + downstream plectoneme sizes from panel (C), which shows an increase during transcription. Solid line represents moving average-filtered data (1 Hz). (**H**) Same as (D), but for N = 21 individual kymographs (grey lines) and their average (black). Transcription-generated supercoils appear to preferentially reside near the ends of the tethered DNA molecule. (**I**) Same as (E), but for the kymographs analyzed in panel (H). (**J**) Same as (F), but for the data analyzed in panel (I), again showing no statistical difference. Scale bars depict 1 µm. Statistical analyses in (F, J) consisted of unpaired, two-tailed t-tests (n.s. = non-significant with p >0.05). See also Supplementary Figures S3, S4, and Supplementary Videos V4, V5.

The plectoneme position distribution (relative to the average RNAP position and the DNA tethering locations) showed a distinct structure (Figure 3D), in contrast to the largely homogeneously distributed plectoneme distribution for bare DNA (Figure 1D). Virtually no plectonemes existed right at the RNAP position (Figure 3D), which is as expected since RNAP is considered a twist-diffusion barrier in the twin-supercoiled domain model. Instead, plectonemes resided up- and downstream of the RNAP. While we observed a high probability of supercoils adjacent to the RNAP (where their formation is facilitated by the bended DNA structure [37], [54], [64], [65]) we noted an even higher probability of supercoils close to the DNA tethering points. The latter observation is consistent with a recent simulation [60] that showed that transcription-induced DNA torsion dissipates much slower (by ~2 orders in magnitude) in writhe (i.e. in plectonemes) than in twist, resulting in plectonemes forming at a distance from the RNAP or upon reaching a twist-diffusion barrier, such as the DNA tether points in our assay.

What is the symmetry of the supercoil density generated up- and downstream during transcription? We found that the fractions of the total plectoneme size were very similar for the up- and downstream supercoils, exhibiting an equal partitioning (0.505±0.09 and 0.495±0.09 (mean±SD), respectively; Figure 3E,F). These distributions were maintained during active transcription, during which the *total* plectoneme size increased from 7 to 10 kbp over 220 s (Figure 3G). The analysis over multiple (N = 21) DNA molecules exhibited that this behavior was conserved: the transcription-induced supercoils were localized with an increased probability at adjacent to the RNAP, but even more so (~2-fold higher probability) near the tethered DNA end locations (Figure 3H), while the supercoils partitioned symmetrically up- and downstream (Figure 3I,J) with close to 50/50% distribution.

Taken together, this first direct visualization of supercoil generation during transcription confirms the characteristics underlying the twin-supercoiled-domain model, namely that the same amounts of supercoils are simultaneously generated up- and downstream.

### Induction of DNA supercoiling by transcription is independent of the RNA transcript

The theoretical framework of the accepted twin-supercoiled domain model indicates that plectonemes are generated because rotation of RNAP during tracking the DNA helical groove is hampered by friction [21], [22], [55]. What provides that friction, in particular in an *in vitro* setting that lacks the numerous factors that interact with the RNAP and its transcript in cells? Current theoretical models indicate that the viscosity of the surrounding medium plays an important role, where the high viscosity of the cytosol would exert sufficient drag on the RNAP and its transcript [19], [58]–[60], while in a low-viscous aqueous buffer, such a viscous drag would be insufficient. We here sought out to probe this concept experimentally.

First, we confirmed the notion of that an RNA transcript can add drag to the RNAP rotation. The most direct evidence of this resulted from observations that upon photo-induced DNA nicking during transcription (Supplementary Figure S5) the supercoil relaxation time was substantially increased to a median value of ~5 s in the presence of the RNAP with its transcript, which was ~9-fold longer as compared to the median relaxation time observed for a supercoiled DNA without a transcribing RNAP [66]).

Subsequently, we extended our fluorescence transcription assay with the addition of RNase A that should digest any present RNA, including the nascent RNA transcript. To do so, we used the same assay as before with a stalled tertiary complex on the 31 kbp DNA construct (Figure 4A). After transcription re-initiation with 1 mM NTPs, we monitored the generation of transcription-induced supercoils for 5 minutes. We then introduced RNase A (in addition to 1 mM NTPs) and monitored the size and positions of the supercoils and RNAP for another 5 minutes (Figure 4A). Based on the classical twin-supercoiled-domain model, one would expect that existing supercoils would relax within seconds (as observed in nicking events) and that no further supercoiling would be induced since with the loss of the transcript the frictional drag exerted by the RNAP alone would be insufficient to induce supercoiling. In contrast to this prediction, however, we observed that supercoils up- and downstream of RNAP continued to be generated after RNase A addition (Figure 4B,C). Apparently, the RNAP was still able to induce supercoils during transcription. Notably, the transcription proceeded at the same rate (Figure 4D,E) and the symmetric partition was maintained (Figure 4F,G). The statistical analysis of N = 24 individual DNA molecules confirmed that all analyzed molecules exhibited both an increase in transcription-induced supercoils (Figure 4H) and that they maintained a symmetric partitioning of supercoils up- and downstream (Figure 4I,J) after RNase A addition. We conclude that the removal of the RNA transcript did not change the ability and characteristics of the RNAP to induce supercoils. Searching for an explanation of the surprising result, we speculated whether R-loops could play a role (Figure 4K). The rationale underlying this hypothesis is that the nascent RNA transcript might anchor onto the DNA via an R-loop and subsequently restrain the ability of RNAP to rotate around the DNA helix [63], [67]. In the absence of an RNA-processing machinery (such as in our *in vitro* assay), R-loops can readily occur, and sequences favoring R-loops are often located at promoter and termination regions [68], [69]. To test for this scenario, we repeated the transcription experiments with the addition of RNase H, which enzymatically resolves RNA:DNA hybrids and R-loops [70], [71]. The analysis of N = 12 individual DNA molecules did, however, not show any effect of RNase H on the transcription-induced supercoiling, as the RNAP in the presence of RNase H was still able to induce supercoils (Figure 4L) that continued to partition symmetrically up- and downstream (Figure 4M).

**Figure 4.**
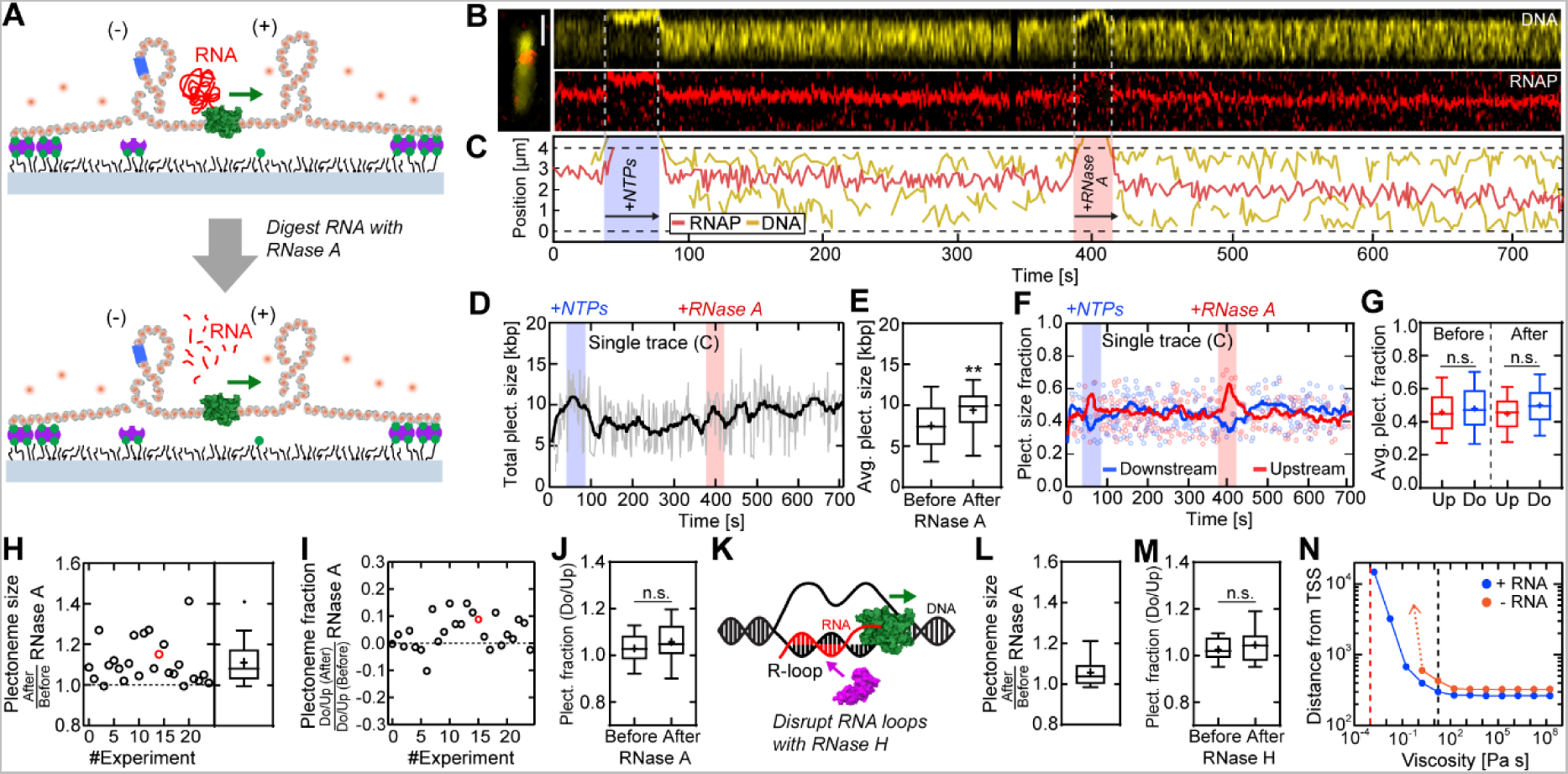
Neither drag of the RNA transcript nor R-loops are essential for generation of DNA supercoils. (**A**) Schematic of the extended transcription assay with addition of RNase A or H. Top: first, transcription occurred on a 31-kbp-long torsionally constrained DNA molecule for 5 minutes. Bottom: then, RNase A was added to digest the RNA transcript, and transcription was continued for another 5 minutes. (**B**) Example fluorescence kymographs of DNA (top, yellow) and RNAP (bottom, red). (**C**) DNA (yellow) and RNAP (red) puncta positions from panel (B) versus time, showing active transcription as well as the generation of up- and downstream supercoils both before and after addition of RNase A. (**D**) Total plectoneme size (i.e., upstream + downstream from panel (C) versus time, showing an increase during transcription, irrespective of RNase A addition. Solid line represents moving average-filtered data to 1 Hz. Blue and red shaded areas depict the time of flushing in NTPs and RNase A, respectively. (**E**) Average total plectoneme size from panel (C) exhibit an increase in supercoil size after the addition of RNase A due to continued transcription. (**F**) Fraction of up- and downstream plectoneme size from panel (C) during transcription versus time. Solid lines depict moving average-filtered data to 1 Hz. (**G**) Average up- and downstream plectoneme size fractions from panel (F), showing no statistical difference before and after adding RNase A. (**H**) Average plectoneme size ratio *‘After/Before RNase A’* (derived like in panel (E)), for N = 24 DNA molecules. The ratio exceeds 1, indicating an increase of supercoiling after the addition of RNase A. Red circle shows the data from panel (C). (**I**, **J**) Ratio of up- and downstream plectoneme sizes for N = 24 DNA molecules (derived like in panel (G)); addition of RNase A did not have any effect. Red circle depicts the data from panel (C). (**K**) Potential R-loops generated during transcription were digested with RNase H using the same experimental assay depicted in panel (A). (**L**) Same as (H), but for RNase H (N = 12). A continuing increase in supercoils is observed after addition of RNase H. (**M**) Same as (J), but for RNase H. No change is observed in up- and downstream plectoneme size fraction ratios after RNase H addition. (**N**) Simulation results of DNA supercoils generated by transcription. Predicted distance from the TSS (Transcription Start Site) at which DNA buckling occurs versus buffer viscosity. Data are taken at a supercoiling density σ = 0.001 in the presence (blue circles) and absence (orange circles) of an RNA transcript. Red dashed line indicates the viscosity of water (1 mPa s); black dashed line indicates the viscosity of bacterial cytoplasm (17.5 Pa s). In the lower left corner (low viscosities and small distances from the TSS), the RNAP tracks the DNA duplex and no supercoiling is induced, while in the upper right corner (high viscosities and large distances from the TSS), the RNAP can no longer track the DNA duplex and supercoiling is induced. In particular, the RNAP without transcript (orange circles) cannot induce sufficient twist at low viscosities (<1 Pa s; orange dotted arrow) to reach the supercoiling density threshold of σ = 0.001. Statistical analyses in (E, G, J, N) consisted of unpaired, two-tailed t-tests (*n.s.* = *non-significant* with p >0.05; ** = p <0.01). See also Supplementary Figures S5, S6.

We thus conclude that neither the RNA transcript nor R-loops play a dominant role in the generation of up- and downstream supercoils. This prompted us to quantitatively estimate the drag, where we used recently published simulations of a theoretical model that accounts for both the mechanical motion of the RNAP and the torsional response of torsionally constrained DNA [72]. Unlike earlier theoretical descriptions of the twin-supercoiled-domain model, this model incorporates experimentally determined mechanical and kinetic properties of DNA, RNAP, and plectoneme formation. We conducted simulations for different buffer viscosities *µ*, ranging from µ = 1.75 μPa s to µ = 17.5 kPa s (for reference: *µ*_water_ = 1 mPa s), in the presence and absence of a nascent RNA transcript (Supplementary Figure S6A,B). As expected, the ability of RNAP to induce supercoils increased with viscosity and RNA transcript length. Considering the viscosity of the bacterial cytoplasm as an example that was previously used in simulations [72] (*µ* = 17.5 Pa s; Figure 4N, black dashed line), the model predicted that the RNAP is able to induce supercoils early on during transcription (i.e. already after ~300 bp), both in the presence and absence of a nascent RNA transcript (Figure 4N; blue and orange circles, respectively), in accordance with previous results [72], [73]. We also simulated the ability of RNAP to induce supercoils using the viscosity of our aqueous buffer solution (*µ* = 1 mPa s; Figure 4N, red dashed line). Here, we observed that RNAP with its transcript was still able to induce supercoils after generating a sizeable transcript (>10 kb; Figure 4N; blue circles). In the absence of an RNA transcript, however, the simulations predicted that the RNAP alone does not induce sufficient twist torque to overcome the DNA buckling energy barrier and induce supercoils in aqueous buffer (Figure 4N; orange circles) – which contradicts our experimental results (Figure 4H-M).

In conclusion, we have used single-molecule assays to provide a direct visualization of the twin-supercoiled domains that are generated by RNAP during transcription up- and downstream. Supercoils were generated in equal portions up- and downstream of the transcribing complex, in full agreement with the classic twin-supercoiled-domain model. Control experiments with RNases A and H revealed that the additional viscous drag of the RNA transcript is *not* necessary for the RNAP to induce supercoils, and accordingly current models cannot fully explain our experimental results that the RNAP ternary complex appears to be able to induce DNA twist on its own. Our results indicate that the hindering of RNAP rotation around the DNA helix is not dominated by the increased frictional drag with the surrounding solvent added by the RNA transcript. An analogous mechanistic question is the apparent ability of individual DNA helicases to induce downstream positive supercoiling during DNA unwinding [74]–[76], for which a theoretical explanation has so far been lacking. Potentially, molecular interactions between the RNAPs/helicases and the DNA may cause additional mechanical friction between the translocating proteins and the DNA at the fork during base pair melting. It will be of interest to verify this hypothesis or other putative causes for the twisting of DNA - independent from the presence of an RNA transcript. Future all-atom molecular dynamics simulations might be able to reveal mechanistic insights to fully describe the twin-supercoiled-domain phenomenon that we here experimentally verified at the single-molecule level.

## DATA AVAILABILITY

The data underlying this article will be shared upon request to the corresponding author.

## SUPPLEMENTARY DATA

Supplementary Data are available online.

## AUTHOR CONTRIBUTIONS

R.J. and C.D. conceived the project. R.J. labelled the *E. Coli* RNA polymerase, performed the experiments with R.B and M.P, and analyzed the data. J.v.T. synthesized the DNA constructs used in this study. R.B. performed the simulations. All authors contributed to writing of this article. C.D. acquired funding and supervised the work.

## ACKNOWLEDGEMENTS

We thank Eli van der Sluis for E. Coli RNA polymerase and σ70 protein purification, and Irina Artsimovitch, Sumitabha Brahmachari, John F. Marko, Herbert Levine, and Erik J. Strobel for discussions.

