## Supplementary Figures for "Single-molecule visualization of twin-supercoiled domains generated during transcription"

### SUPPLEMENTARY DATA

**SUPPLEMENTARY FIGURES**

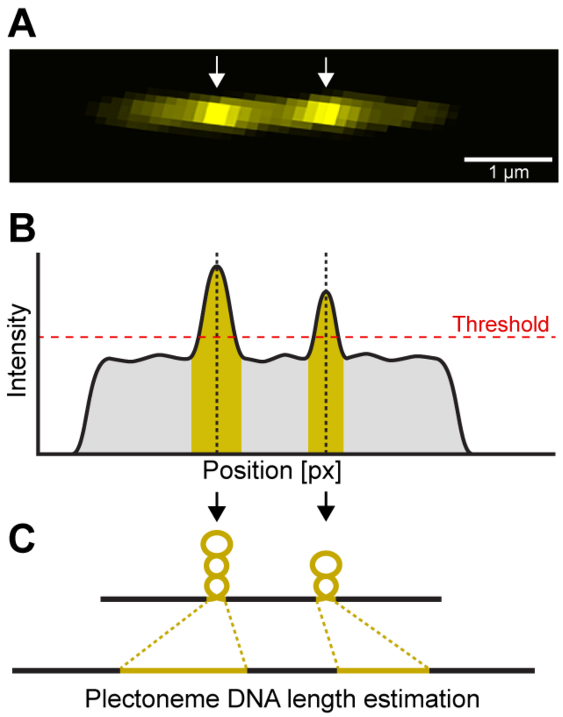


**Supplementary Figure S1. Estimation of plectoneme position and size (in bp) from DNA fluorescence emission intensities.** (**A**) Representative fluorescence emission image of a supercoiled torsionally constrained DNA molecule with two plectonemes (arrows). (**B**) Fluorescence intensity profile of the supercoiled DNA with two plectonemes (position shown by dotted lines) from panel (A). (**C**) The size of the plectoneme is computed as the fraction of intensity attributed to the plectonemes divided by the total intensity along the entire DNA molecule, times the total DNA length in bp. **Related to Figure 1.**

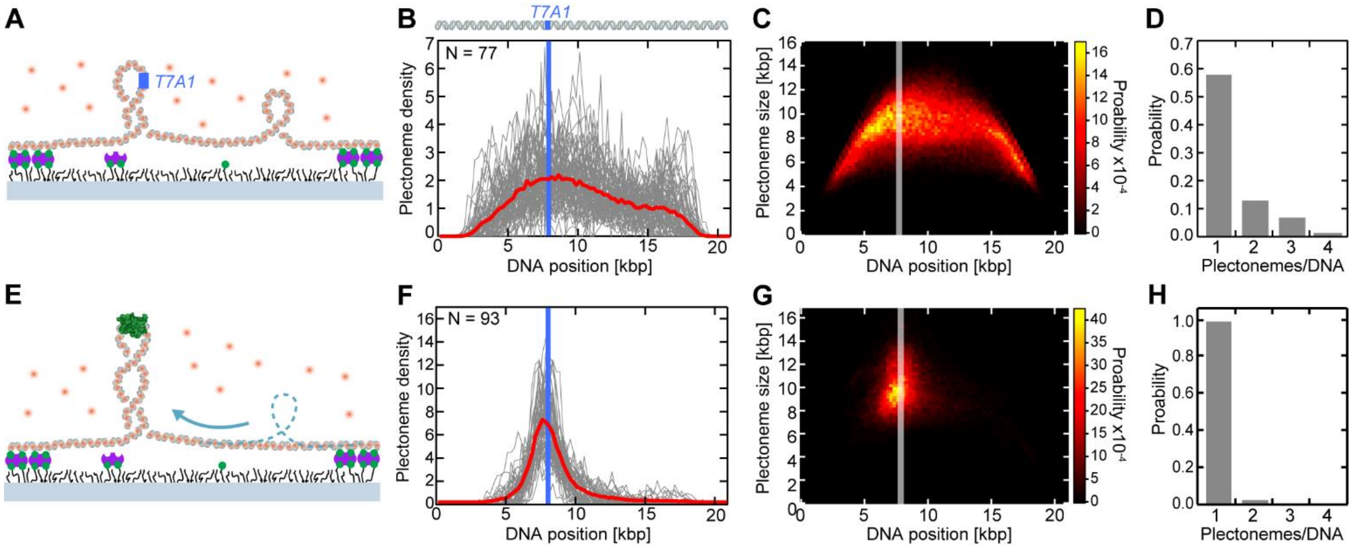


**Supplementary Figure S2. Positive DNA supercoils are merged into large plectoneme pinned by bound RNAP.** (**A**) Single-molecule assay schematic. (**B**) DNA position-dependent positively coiled plectoneme densities for N = 77 individual DNA molecules (grey lines) and their average (red line). The position of the T7A1 promoter is depicted by the blue line. (**C**) DNA position-dependent plectoneme size probability distribution from the data in panel (A). The T7A1 promoter position is represented by the grey line. (**D**) Probability distribution of the number of positively coiled plectonemes co-existing during each frame for DNA molecules measured in panel (A). (**E**) Schematic of the assay with bound RNAP. (**F**) Same as (B), but in presence of bound RNAP for N = 93 individual DNA molecules. (**G**) Same as (C), but in presence of bound RNAP. (**H**) Same as (D), but in presence of bound RNAP, showing only one existing plectoneme upon binding RNAP. **Related to Figures 1, 2.**

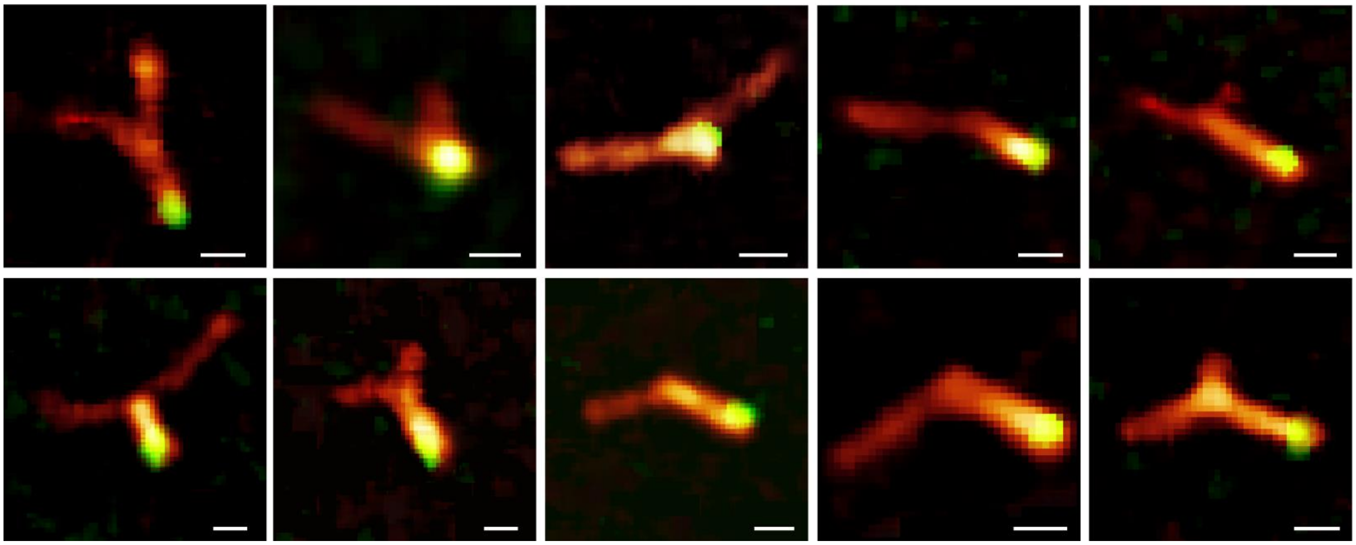

**Supplementary Figure 3. Bound RNAP resides at the apex of pinned DNA plectonemes.** Further dual-color fluorescence images from side-flow experiments shown in Figure 2H, demonstrating that the RNAP (green) positions at the apex of pinned DNA (orange) plectonemes. Scale bars depict 1 µm. **Related to Figure 2.**

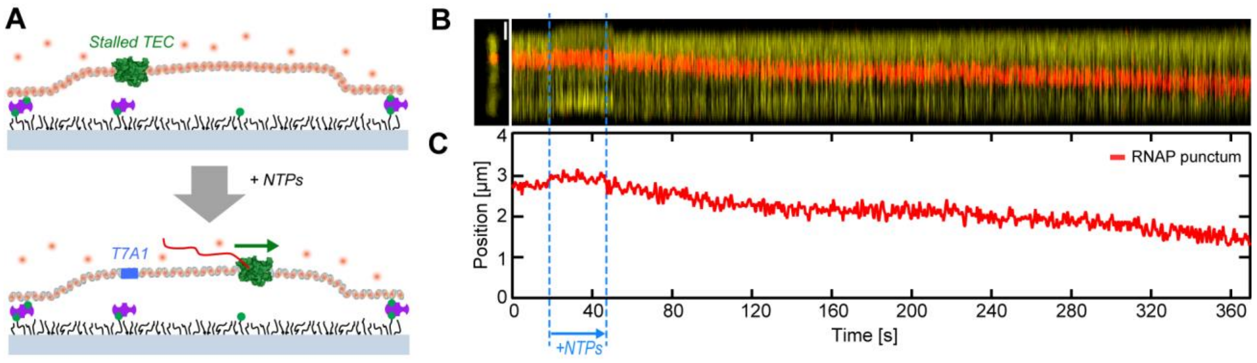


**Supplementary Figure S4. Transcription on torsionally unconstrained linear DNA.** (**A**) Schematic of the single-molecule transcription assay similar to Figure 3A, but for nicked linear DNA molecules. (**B**) Superimposed example kymograph of DNA (yellow) and RNAP (red). No supercoils are being generated after transcription restart *via* NTP addition. (**C**) RNAP punctum position on the DNA from panel B versus time after transcription restart via NTP addition. **Related to Figure 3.**

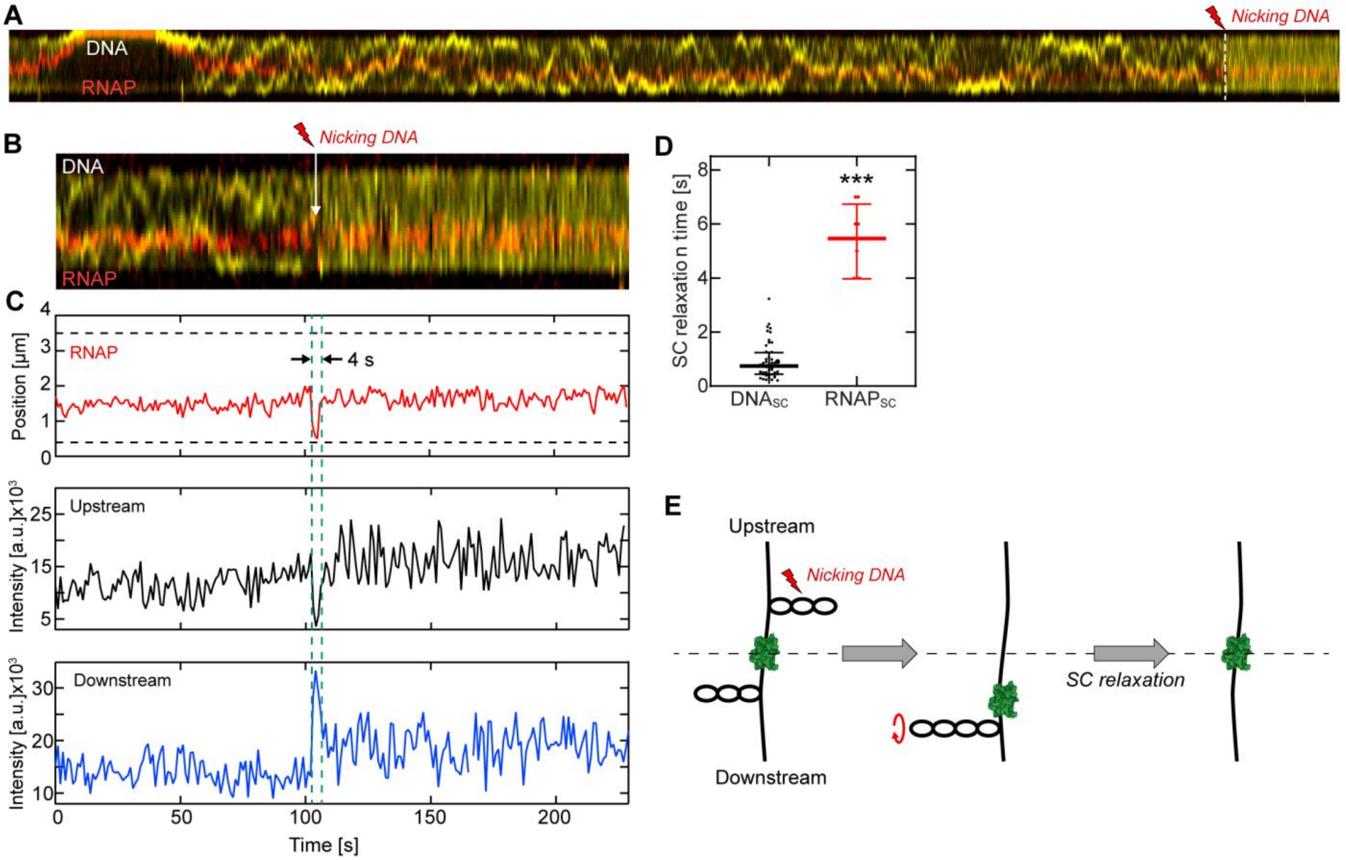


**Supplementary Figure S5. Supercoil relaxation slowed by the rotational drag added by RNAP and transcript.** (**A**) Example fluorescence kymograph of DNA (yellow) and RNAP (red) with a DNA nicking event (lightning symbol, red). Frame rate was 1 Hz. (**B**) Magnified region of panel (A) around the DNA nicking event. (**C**) (Top) RNAP position exhibits a sudden ~1.5 µm movement into the RNAP downstream direction (i.e. down in y-axis), followed by a recovery to its initial position after ~4 s (between cyan dashed lines). The horizontal black dashed lines represent the tethered end points of the DNA molecule. (Center) The fluorescence intensity of the DNA upstream RNAP exhibits a sudden and strong (~3-fold) decrease. (Bottom) The downstream intensity simultaneously shows an abrupt increase of similar magnitude, both occurring at the same time when the RNAP started to move and with a comparable recovery time of ~4 s. (**D**) The time necessary to relax supercoils after DNA nicking of bare $DNA_{SC}$ (left, black, data from reference [66]) is ~9-fold faster than the supercoil recovery during transcription ($RNAP_{SC}$) that is measured upon nicking the DNA at one side of the twin domain (median 0.61 s vs 0.54 s, respectively. Plot shows individual supercoil relaxation times and median±1.5 IQR. (**E**) Schematic showing fast supercoil relaxation at the DNA upstream of RNAP where nicking occurred, which caused the RNAP to move down due to the sudden loss of twist-induced DNA tension upstream. Relaxation of the supercoiled DNA downstream of RNAP occurred slower, presumably due to the additional rotational drag of the bound RNAP and its transcript. After complete supercoil relaxation and tension equilibrium, the RNAP is located again at the same position as before DNA nicking. Statistical analysis in panel (D) consisted of an unpaired, two-tailed t-test (*** = $p < 0.001$). **Related to Figure 4.**

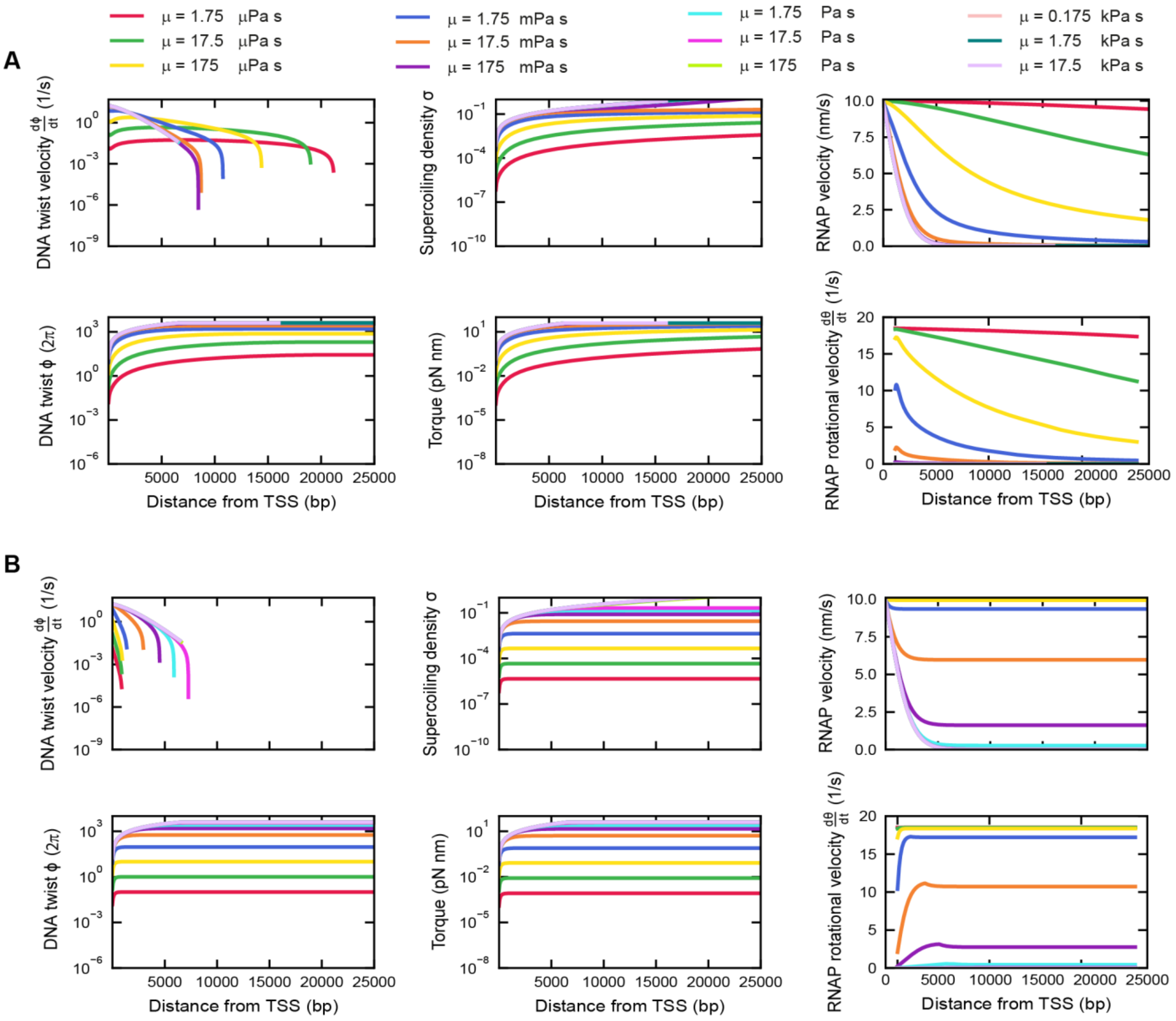


**Supplementary Figure S6. Simulation of supercoil generation in dependence of viscosity and transcript length.** (**A**) From top left to bottom right: calculated DNA twist velocity, accumulated downstream DNA supercoiling density σ, linear RNAP transcription velocity, accumulated DNA twist downstream of RNAP, generated torque, and RNAP rotational velocity around DNA – for an RNAP transcribing up to 25 kbp at different solution viscosities μ, ranging from μ = 1.75 μPa s to μ = 17.5 kPa s, covering the range including the approximate viscosity of water (μ = 1 mPa s; cf. the dark blue line) and the bacterial cytoplasm (μ = 17.5 Pa s; pink line). In these calculations, no external DNA tension is applied (force = 0 pN). The RNAP transcription velocity is measured as the linear velocity along the DNA. The RNAP can rotate around DNA in solutions with low viscosity (bottom right panel), inducing only a marginal twist in the DNA (left panels). RNAP rotation is inhibited at large viscosities due to the frictional drag. The increase in DNA torsional tension with distance from the TSS site (Transcription Start Site) slows down the RNAP transcription velocity (top right panel). However, the DNA twist velocity approaches zero during transcription of the 25 kbp gene (top left panel) at which the RNAP has to rotate in order to further transcribe the gene, albeit slowly. The translocation of the RNAP along the gene in between fixed ends causes the supercoiling density of the downstream segment and the torque to further increase as the downstream segment becomes shorter (middle panels). (**B**) As (A), but without taking into account the RNA transcript that adds to an increase of rotational drag. **Related to Figure 4.**

**SUPPLEMENTARY VIDEOS**

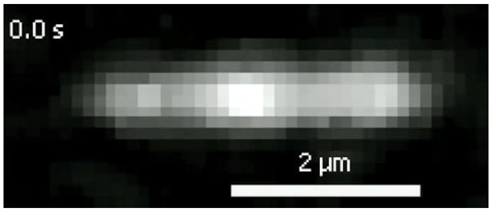


**Supplementary Video V1.** Representative real-time fluorescence video that captures the behavior of a Sytox Orange-induced negatively supercoiled DNA that is attached to a surface in absence of external flow. The video features dynamic characteristic bright spots indicative of supercoiled DNA in form of plectonemes that can nucleate, diffuse along the DNA, and terminate over time. **Related to Figure 1.**

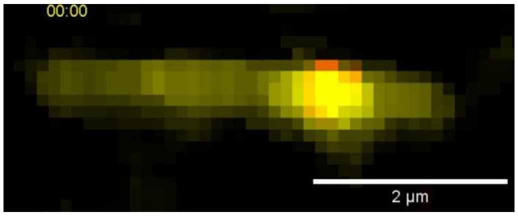


**Supplementary Video V2.** Real-time dual-color fluorescence video showing negative DNA supercoils (yellow) that are merged into a large plectoneme that is pinned by bound RNAP (red). **Related to Figure 2.**

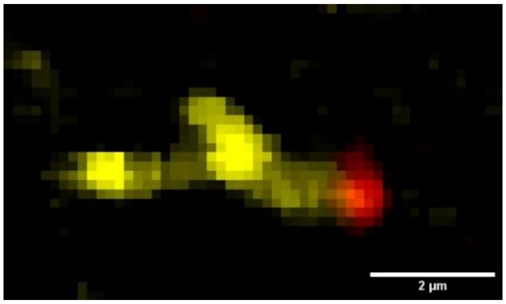


**Supplementary Video V3.** Real-time dual-color fluorescence video performed with side flow (from left) showing a long DNA plectoneme (yellow) pinned by RNAP (red) that resides at the apex of the plectoneme. **Related to Figure 2.**

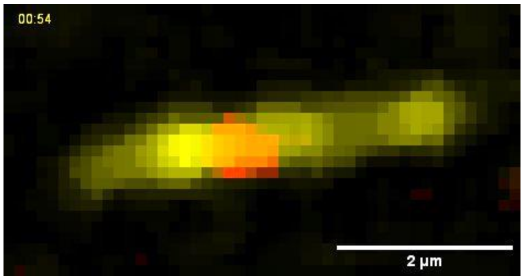


**Supplementary Video V4.** Real-time dual-color fluorescence video of RNAP (red) transcribing a torsionally constrained DNA construct (yellow). Bright spots are indicatives of transcription-generated plectonemes that diffuse along the DNA. **Related to Figures 3, 4.**

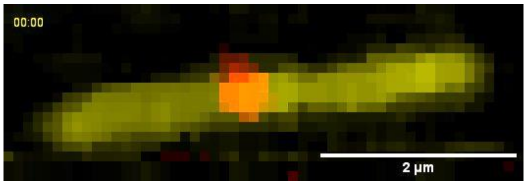


**Supplementary Video V5.** Real-time dual-color fluorescence video of RNAP (red) transcribing a linear, non-torsionally constrained DNA construct (yellow). **Related to Figure 3.**
